# A novel *adipose* loss-of-function mutant in *Drosophila*

**DOI:** 10.1101/2023.08.18.553914

**Authors:** Nicole A. Losurdo, Adriana Bibo, Jacob Bedke, Nichole Link

**Author notes:** Corresponding author: N.L. The Solomon H. Snyder Department of Neuroscience, Johns Hopkins University, Baltimore, MD, 21218, USA.

## Abstract

To identify genes required for brain growth, we took an RNAi knockdown reverse genetic approach in Drosophila. One potential candidate isolated from this effort is the anti-lipogenic gene *adipose* (*adp*). *Adp* has an established role in the negative regulation of lipogenesis in the fat body of the fly and adipose tissue in mammals. While fat is key to proper development in general, *adp* has not been investigated during brain development. Here we found that RNAi knockdown of *adp* in neuronal stem cells and neurons results in reduced brain lobe volume and sought to replicate this with a mutant fly. We generated a novel *adp* mutant that acts as a loss-of-function mutant based on buoyancy assay results. We found that despite a change in fat content in the body overall and an increase in the number of larger (>5μm) brain lipid droplets, there was no change in the brain lobe volume of mutant larvae. Overall, our work describes a novel *adp* mutant that can functionally replace the long-standing *adp*^*60*^ mutant and shows that the *adp* gene has no obvious involvement in brain growth.

## INTRODUCTION

Many genes associated with human disease were discovered and studied through model organisms, including genes required for brain growth^1–4^. Human variants in critical neurodevelopmental genes can cause microcephaly, which is a rare neurodevelopmental disorder characterized by a reduced occipital frontal circumference (OFC) of two standard deviations (SD) or more below the mean for a child’s age and sex^5–7^. Half of all known causes of microcephaly are from genetic mutations, suggesting that additional cases of genetic microcephaly might be found by assessing genes for neurodevelopment defects in model organisms^3,4,6–8^. We hypothesized that by using a reverse genetics approach, we would isolate novel genes required for neurodevelopment and linked to human disease. We used commercially available *Drosophila* RNAi lines to knock down candidate genes in developing brains and screened third-instar larval *Drosophila* for brain size differences. Through this screen, we identified a potentially novel candidate gene in neurodevelopment, *adipose* (*adp*).

*Adipose* was first characterized in the 1950s when a wild strain of fly, *adp*^*60*^, was isolated in Africa^9^. These *adp*^*60*^ flies had a 23-base pair deletion in the middle of the *adp* gene, leading to increased lipogenesis^10–12^. Adult *adp*^*60*^ flies were resistant to starvation compared to Oregon R controls, and both the larvae and adults were shown to have higher triglyceride levels^10–12^. Further work validated that *adp* functions as an inhibitor of lipogenesis, and the *adp*^*60*^ flies carried a loss of function allele, demonstrating an increase in fat storage. While the lipogenesis phenotypes of *adp*^*60*^ have been well characterized, it is currently unknown whether *adp* plays a role in neurodevelopment.

The importance of fat during brain development has been well demonstrated across species. In vertebrates, fat is necessary to produce the myelin sheath covering axons. In the developing fly brain, neural stem cells, called neuroblasts, reside in stem cell niches protected by lipid droplets^13,14^. No work has shown whether *adp* is necessary to produce brain lipid droplets in larval *Drosophila*, but we hypothesized that *adp* is necessary for neurodevelopment and is likely to function through lipid droplet production in the brain. In this paper, we generated a novel *adp* mutant that behaves as a loss of function mutant that replicates high-fat content as previously shown. We also demonstrate that *adp* is not necessary for neurodevelopment in the fly nor for lipid droplet production in the brain.

## RESULTS

In an effort to identify conserved pathways required for brain growth and novel players in neurodevelopment, we screened a collection of *Drosophila* genes for function in the brain using *in vivo* RNAi. We crossed *UAS-RNAi* flies to either *inscuteable-GAL4* (*insc-GAL4*)^15^ or *neuronal synaptobrevin-GAL4* (*nsyb-GAl4*) to knock down genes of interest in neuronal stem cells or postmitotic neurons, respectively. Developing brains from late third-instar larvae were collected for volumetric analysis to assess brain lobe growth^8,16^. Knockdown of *adipose* (*adp*) in both neuronal stem cells and post-mitotic neurons resulted in significantly reduced brain lobe volume compared to the control knockdown of GFP (Figure 1A-F). These results indicate that *adp* may be involved in neurodevelopment.

**Figure 1:**
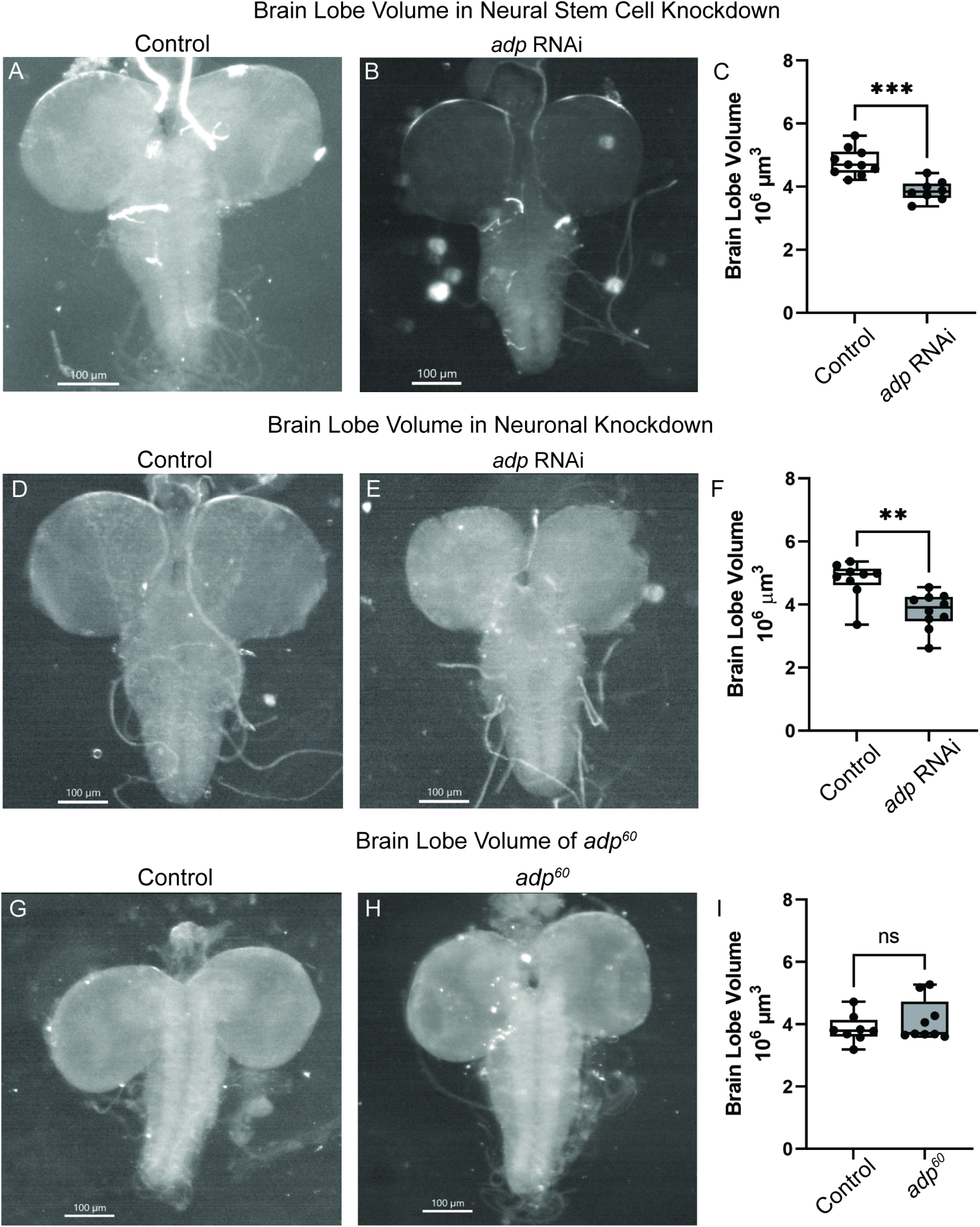
RNAi knockdown of *adp* but not the *adp*^*60*^ mutant results in significantly reduced brain lobe volume. Bright-field images of third instar larval brains just prior to pupation with (A, D) GFP knockdown (VALIUM22-EGFP.shRNA.1) and (B, E) *adp* knockdown (TRiP.HMC006600) in neural stem cells (A-C, *insc-GAL4*) or post-mitotic neurons (D-F, *nsyb-GAL4*). (C) Brain lobe volume of neural stem cell knockdown of GFP (Control) or *adp* in third instar larvae; each dot represents one brain lobe (n=8-10). Knockdown of *adp* in neural stem cells results in significantly reduced brain lobe volume compared to control (independent t-test, t=4.932, df=16, p=0.0002). (F) Brain lobe volume of post-mitotic neuronal knockdown of GFP (Control) or *adp* in third instar larvae, each dot represents one brain lobe (n=9-10). Knockdown of *adp* in post-mitotic neurons results in significantly reduced brain lobe volume compared to control (independent t-test, t=3.694, df=17, p=0.0018). Bright-field images of third instar larval brains just prior to pupation of (G) Oregon-R and (H) *adp*^60^ fly stocks. (I) Brain lobe volume of Oregon-R (Control) or *adp*^*60*^ third instar larvae, each dot represents one brain lobe (n=8-9). The *adp*^*60*^ mutant does not differ in brain lobe volume compared to an *Oregon-R* control (independent t-test, t=0.9658, df=15, p=0.3495). Sequencing of *adp*^*60*^ showed no mutation in the *adp* gene.

To verify that reduced brain lobe volume resulted from a loss of *adp* and not off-target RNAi effects, we assessed a known loss-of-function mutation called *adp*^*60* 9,10^. Adult male *adp*^*60*^ flies are resistant to starvation, and both adult males and third-instar larvae have increased triglyceride content compared to wild-type animals, indicating that Adp functions to inhibit lipogenesis^10–12^. However, third-instar larval *adp*^*60*^ brains were not significantly different in size compared to *Oregon-R* control brains (Figure 1G-I). This negative result could mean that *adp* is either unnecessary for brain development or the mutation in *adp*^*60*^ may no longer be present.

We aimed to verify the mutation by sequencing the *adp*^*60*^ fly with the same primers described in the initial characterization^11^. The published mutant contains a 23-base pair deletion that removes nucleotides 1153 to 1176 in exon 2, resulting in an early stop codon in the predicted protein. The *adp* sequence in available *adp*^*60*^ mutants matches the reference genome with no deletion present in both the main and backup *adp*^*60*^ Bloomington stocks, indicating that *adp*^*60*^ is no longer a mutant allele of *adp*.

Due to the lack of an available *adp* mutant, we sought to generate our own loss-of-function mutant using the TRiP-CRISPR toolbox^17–27^. We crossed flies expressing Cas9 in the germline (*nanos-Cas9*) together with flies expressing single guide RNA for *adp* targeting base pairs 221-243 in the first exon. F1 males are expected to have germline mutations in *adp*, and F2 founder flies were isolated to generate 10 independent mutant *adp* lines. Initially, three lines were genetically characterized using PCR and Sanger sequencing of the *adp* locus. Surprisingly, all three contained small INDELs in the guide RNA target region, resulting in early stop codons in the predicted protein. The remaining 7 lines have not been sequenced. We decided to focus on a single mutant, which we named *adp*^*1*^. The *adp*^*1*^ mutant contains a frameshift mutation (c.237_238insT) predicted to result in an early stop codon in the first exon (p.Asp134Ter) (Figure 2B). The *adp*^*1*^ mutant is homozygous fertile and viable. Due to how early the predicted truncation mutation appears, we predict *adp*^*1*^ to be a loss-of-function mutation.

**Figure 2:**
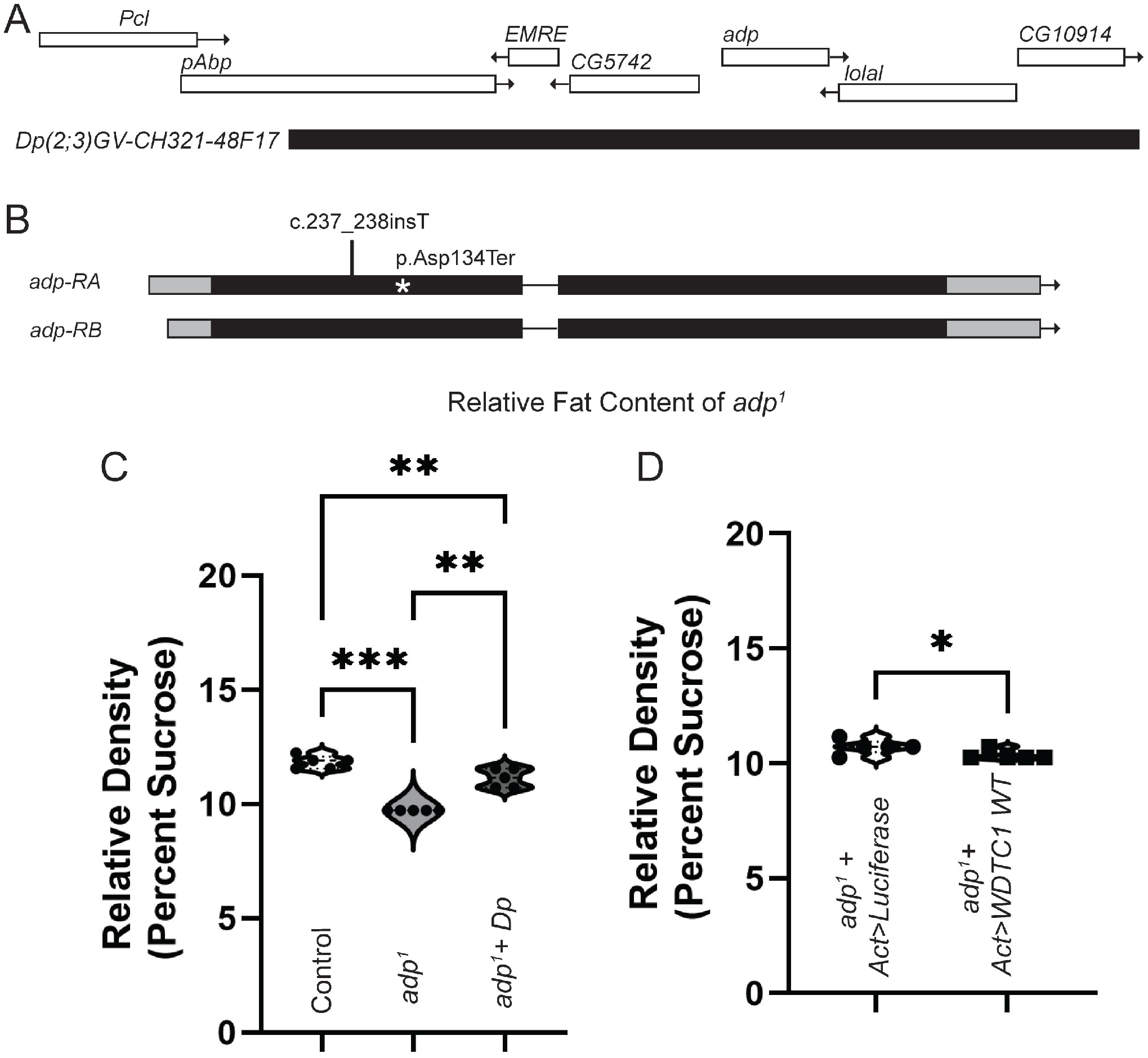
*adp*^*1*^ has higher fat content than controls. (A) Genomic region of *adp* and location of duplication line used for rescue. (B) Sequencing of fly *adp*^*1*^ showed a frameshift mutation (c.237_238insT) resulting in an early stop codon p.Asp134Ter marked with the asterisk. (C) Buoyancy assay comparing control (*white berlin*), *adp*^*1*^, and *adp*^*1*^ plus the genomic duplication noted in (A), each dot represents a replicate experiment of around 20-30 larvae per genotype (total n sizes from all replicates: control n=100, mutant n=139, rescue n=125). Loss of function *adp*^*1*^ larvae float at a lower density than control larvae, while a genomic duplication of fly *adp* is able to rescue (repeated measures one-way ANOVA, F=117.9, dF=14, p=0.0001, with Tukey’s multiple comparisons control vs. *adp*^*1*^ p=0.0002, control vs *adp*^*1*^ plus duplication p=0.0033, and *adp*^*1*^ vs *adp*^*1*^ plus duplication p=0.0035). (D) Buoyancy assay comparing *adp*^*1*^ plus ubiquitous expression of Luciferase to control for GAL4 expression and *adp*^*1*^ plus ubiquitous expression of the WT human ortholog, *WDTC1*, each dot represents a replicate experiment of around 20-30 larvae per genotype (total n sizes from all replicates: Luciferase n=149, *WDTC1* n=140). Ubiquitous expression of *WDTC1* does not rescue *adp*^*1*^ buoyancy phenotype (paired t-test, t=3.979, dF=4, p=0.0164).

Previous research shows that loss of *adp* results in increased fat stores, so we verified this observation with our new mutation before assessing our mutants for neurodevelopmental phenotypes^9–12^. Using a buoyancy assay, we evaluated changes in fat storage at the third-instar larval stage^28^. *adp*^*1*^ mutant and *white berlin* control third-instar larvae were floated in a sucrose solution where density was increased until all larvae were floating. The density at which all larvae floated was recorded and analyzed. As expected, *adp*^*1*^ mutants float at a lower density than *white berlin* controls, indicating an increase in fat content compared to controls (Figure 2C). To show that the increase in fat storage was due to mutations in *adp* and not background variability or off-target effects of CRISPR mutagenesis, we introduced a 80kb genomic duplication line containing the *adp* locus and again tested buoyancy. We were able to significantly rescue the fat phenotype, indicating that loss of *adp* causes increased fat stores (Figure 2C).

The human ortholog of *adp, WDTC1*, has also been shown to inhibit lipogenesis, and mutants of *WDTC1* lead to increased triglyceride content in mice, similar to phenotypes we observe in flies^12,29–31^. Therefore, we wanted to assess if *WDTC1* can functionally rescue our *adp*^*1*^ mutant’s buoyancy phenotype. Ubiquitous expression of *WDTC1* cDNA in our *adp*^*1*^ background could not rescue the buoyancy phenotype observed in our *adp*^*1*^ mutant (Figure 2D). The human and fly genes may not be functionally conserved or work through different molecular mechanisms despite both being shown to regulate similar processes.

Having confirmed that *adp*^*1*^ displays similar loss of function phenotypes based on previous research, we wanted to know if *adp* loss results in brain growth perturbations to validate our RNAi data. The larval brain contains lipid droplets that help maintain the stem cell niche, so we first assessed whether *adp*^*1*^ has changes in lipid droplet number that would indicate changes in lipogenesis in the brain^13,14^. We performed Nile Red staining in third instar larval brains and quantified the total number of lipid droplets in *adp*^*1*^ and *white berlin* brains^13,32^. While *adp*^*1*^ mutants display no significant difference in the total number of lipid droplets per micrometer cubed, they do have significantly more droplets over 5μm in diameter (Figure 3A-D). This suggests that there is an increase in lipogenesis in the brain, and *adp* may be involved in controlling lipid droplet size.

**Figure 3:**
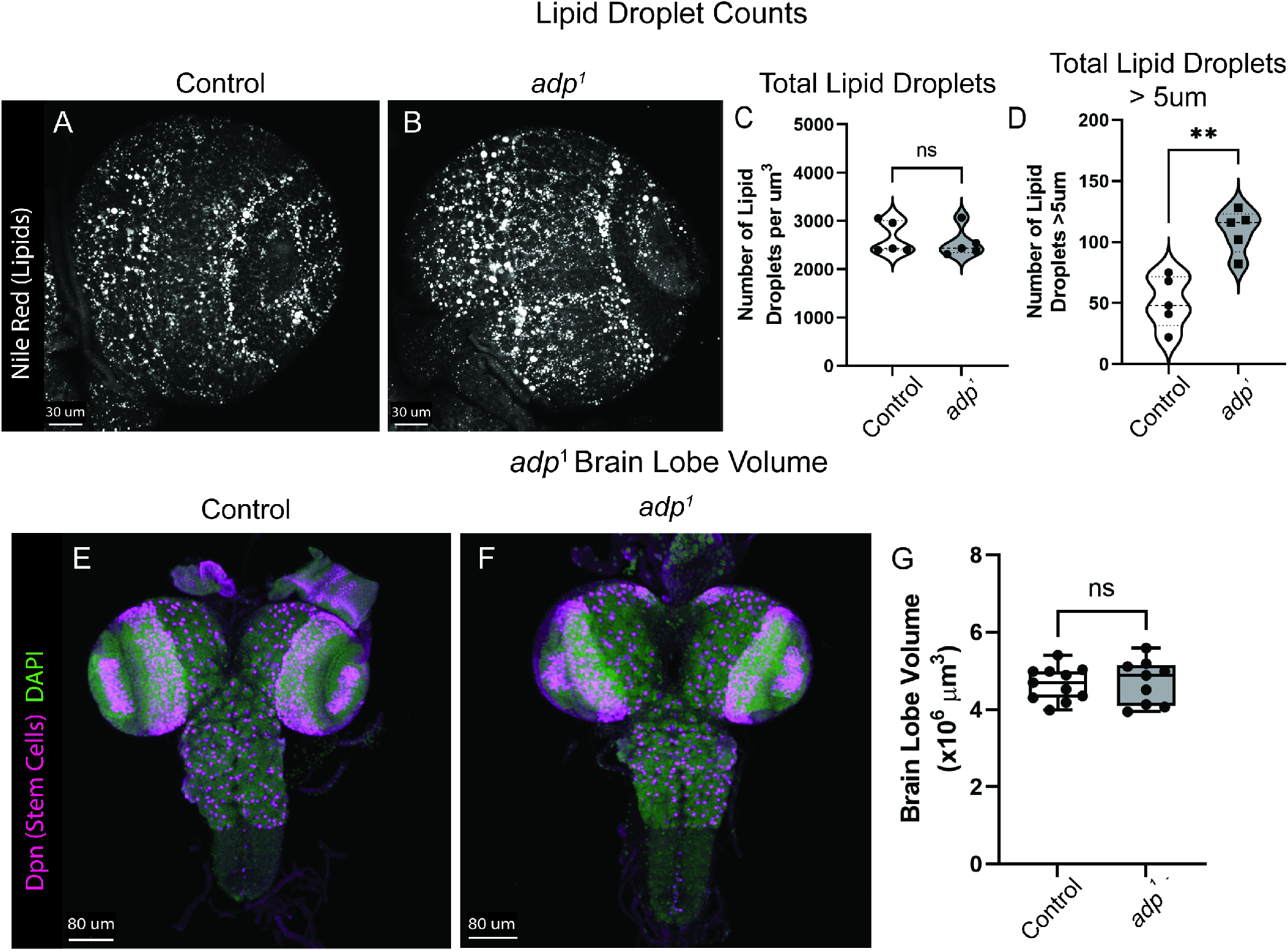
*adp*^*1*^ shows no difference in number of brain lipid droplets nor brain lobe volume. Nile red staining in the third-instar brain lobe showed no difference between (A) controls (*white berlin*) and (B) *adp*^*1*^ in the total number of lipid droplets, quantified in (C), where each dot represents a single brain lobe (independent t-test, t=0.4996, dF=8, p=0.6308). (D) *adp*^*1*^ mutants have significantly more lipid droplets greater than 5μm in diameter, where each dot represents a single brain lobe (independent t-test, t=4.703, dF=8, p=0.0015). Immunohistochemistry staining of third-instar larval (E) *white berlin* control and (F) *adp*^*1*^ brains showing Deadpan (Dpn, stem cells, magenta) and DAPI (DNA, green). (G) Brain lobe volume of white berlin (Control) or *adp*^*1*^ third instar larvae, each dot represents one brain lobe (n=9-11). The *adp*^*1*^ mutant does not differ in brain lobe volume compared to a *white berlin* control (independent t-test, t=0.2020, df=18, p=0.8422).

Finally, we wanted to determine if *adp* loss of function affected brain growth to replicate our RNAi data in a loss-of-function model (see Reagent Table for specific antibodies used). We quantified the brain lobe volume of *adp*^*1*^ and *white berlin* third instar larvae but found no significant difference, indicating that *adp* is unnecessary for brain growth (Figure 3E-G). Our results also suggest that the initial RNAi results we found may have been due to off-target effects or as a result of cell specific knockdown.

## DISCUSSION

In this study, we generated a novel *adp* mutant to investigate its role in neurodevelopment. We showed that the previous standard *adp* mutant available from the Bloomington Stock center, *adp*^*60*^, did not carry the described mutation. Our new mutant, *adp*^*1*^, acts as a loss of function exhibiting the expected phenotype of increased fat stores but failed to show any changes in neurodevelopment.

Previous discovery and description of the *adp*^*60*^ fly demonstrated *adp’s* role in the negative regulation of lipogenesis^9–12^. Adult *adp*^*60*^ flies had increased survival in starvation scenarios, and both larvae and adults exhibited increased triglyceride storage in the fat body. This research validated the use of *adp*^*60*^ as a negative control for lipogenesis research^28^. However, our sequencing of the *adp*^*60*^ stock obtained from the Bloomington Stock Center using the primers described in the original sequencing showed sequence that was identical to the reference genome^11^. The 23 base pair deletion was no longer present in this stock, making it wild-type. In order to perform our own tests of the role of *adp* in neurodevelopment, we generated a new loss of function mutant.

Our *adp*^*1*^ mutant larvae float at a lower density than controls, equating to a higher fat-to-muscle ratio, consistent with an *adp* loss of function phenotype. Despite the previous literature on *adp*^*60*^ not specifically quantifying the buoyancy of *adp* mutant larvae, most buoyancy protocols suggest using *adp*^*60*^ larvae as a control for higher fat content^28^ (Figure 2C). These results validated that our newly generated mutant acts as a loss of function and demonstrated that our mutant could functionally replace *adp*^*60*^ as a control in lipogenesis research.

While *adp* has not previously been linked with neurodevelopment, expression data show that transcript and protein are present in the third instar larval central nervous system^33–36^. This expression profile and our RNAi knockdown results (Figure 1A-F) indicated that *adp* might have a role in brain development. Lipid droplets provide a hypoxic environment for neuroblasts to proliferate efficiently and protect the neuroblasts from assault by reactive oxygen species^13,14^. Disruption of lipid droplet production in glia reduces neuroblast proliferation in hypoxic conditions^13^. Adp inhibits triglyceride storage but has not previously been linked to lipid droplet production in the larval brain. Here we show that *adp* loss of function does not affect the total number of lipid droplet in the third-instar larval brain but increases the number of droplets greater than 5μm in diameter (Figure 3A-D), indicating that *adp* may regulate lipid droplet size. The vertebrate ortholog of *adp, WDTC1*, suppresses lipogenesis through histone modification, but this molecular function has yet to be investigated in the fly^31^.

Despite our RNAi data indicating that *adp* may function in the developing brain, we failed to see a difference in brain size (Figure 3E-G). *Adp*^*1*^ acts as a loss of function mutant based on its mutation, fat phenotypes, and ability to rescue with duplication, so the lack of brain size phenotype confirms *adp* is not necessary for proper brain development. RNAi can have off-target effects, which is hypothesized here. We are confident that *adp* is not a vital component of brain volume regulation.

## Data Availability Statement

Strains and plasmids are available upon request. The authors affirm that all data necessary for confirming the conclusions of the article are present within the article, figures, and tables.

## ACKNOWLEDGMENTS

Stocks obtained from the Bloomington Drosophila Stock Center (NIH P40OD018537) were used in this study. Human ORFs used in this study were provided by the University of Utah HSC Core Research Facilities as part of a partnership with the Huntsman Cancer Institute and individual contributing investigators. Thank you to members of the Link lab for thorough reading of the manuscript. We acknowledge the information provided by FlyBase using release FB2023_04^37^.

## METHODS

### Fly Lines

The following fly lines were used: *adp* RNAi (*P{TRiP*.*HMC06600}attP40*), EGFP RNAi (*P{VALIUM22-EGFP*.*shRNA*.*1}attP40*), *inscuteable-GAL4* (*P{w[+mW*.*hs]=GawB}insc[Mz1407])*^15^, *neuronal Synaptobrevin-GAL4* (*P{y[+t7*.*7] w[+mC]=nSyb-GAL4*.*P}attP2*), *adp*^*60* 9,10^, *Oregon-R* ^9,10^, *nanos-Cas9* (*P{y[+t7*.*7] v[+t1*.*8]=nos-Cas9*.*R}attP2*)^17^, *adp* snRNA:U6:96Ac (*P{y[+t7*.*7]*

*v[+t1*.*8]=TKO*.*GS04840}attP40*)^17–27^, *adp*^*1*^(this study), *white berlin* (*w*^*1118*^) ^38^, *(P{UASt-CD8-GFP}*)^39^, *actin-GAL4* (*P{Act5C-GAL4} 17bFO1*)^26^, *P{UASt-WDTC1}VK33* (this study), *w[1118]; Dp(2;3)GV-CH321-48F17, PBac{y[+mDint2] w[+mC]=GV-CH321-48F17}VK00031*. All flies were maintained at 25°C and grown on Archon glucose formula medium in plastic vials. Crosses were performed at the temperature indicated (18°C, 25°C, or 29°C). Brain volume measurements were conducted in late wandering 3rd instar larvae identified by gut clearance and extruding spiracles^8,16^.

### RNAi Knockdown of *adp*

Male *adp* RNAi and EGFP RNAi flies were crossed with either *insc-GAL4* or *nsyb-GAL4* females for knockdown in neural stem cells or post-mitotic neurons, respectively. Crosses were set at 29°C. Third-instar larvae were selected for brain lobe volume analysis.

### Immunohistochemistry for Brain Volume

Late third instar larval brains were dissected and immediately fixed in 4% paraformaldehyde in Phosphate Buffered Saline + 0.3% TritonX (PBST) for 20 minutes^8,16^. Brains were washed with PBST three times for five minutes and blocked twice with PBST + 1% w/v Bovine Serum Albumin (PBSTB) for 30 minutes before blocking with PBSTB + 5% Normal Donkey Serum (NDS) for 30 minutes.

Brains were incubated with 1:1000 rat anti-Deadpan (neuroblast marker, Abcam, ab195173) in PBSTB overnight at 4°C. Primary antibody was removed from brains before washing three times with PBSTB for 20 minutes. Next, the brains are incubated with secondary antibodies 1:500 Donkey anti-Rat fluorophore 647 (Jackson ImmunoResearch Laboratories Inc., 712-605-153) and 1:1000 DAPI for one hour at room temperature. Finally, brains were washed with PBST four times for 10 minutes before being mounted for confocal microscopy.

### Generation of CRISPR Mutants

The *adp*^*1*^ mutant was generated from the TRiP-CRISPR stocks and TRiP-CRISPR Knockout (TRiP-KO) protocol^17–27^. Ten *nanos-Cas9* females were crossed with 6 *adp* sgRNA males at 25°C to generate germline mutations in *adp*^17–20,23,26^. Fifteen F1 male flies (*y*,*v*,*sc*,*sev; adp sgRNA/+; nanos-Cas9/+*) were then crossed with 15 *y*,*v*,*sc*,*sev; lethal/CyO* females to isolate mutant animals. Both *nanos-Cas9* and *adp* sgRNA constructs are tagged with *y*^*+*^,*v*^*+*^. F2 individuals were selected for *y*^*-*^ *and v*^*--*^ to ensure the removal of the Cas9 and sgRNA sequences and balance *adp* mutations. Once F3 larvae appeared, the founder F2 individual was removed from the tube for sequencing. Ten mutant stocks were established using this method. All work in the rest of this paper was performed with the *adp*^*1*^ mutant.

### Sequencing and Primers

Founder F2 adults were squished with fresh squishing buffer (10 mM Tris pH 8.2, 25 mM NaCl, 1 mM EDTA, and 200 ug/mL Proteinase K). The lysate was incubated for 30 minutes at 37°C degrees and then for 10 minutes at 85°C. 2 uL of the lysate was used for sequencing. The following primers were used for PCR: 5’-AACAAGTGTCATAATCCTATCCACAGCA-3’ and 5’-

TGCATGCAGCCAATATAGATCAAGATG-3’. PCR products were purified and sequenced using the same primers. Sequencing of *adp*^*1*^ showed a single insertion c.237_238insT resulting in an early stop codon p.Asp134Ter.

### Buoyancy Assay

To indirectly test the fat content of *adp*^*1*^, we performed a buoyancy assay^28^. Approximately 20-40 late third instar larvae were collected from either *adp*^*1*^ or *white berlin* vials and placed in 50 mL conical tubes with a starting solution of 11.5 mL PBS and 9 mL 20% w/v Sucrose in PBS. Samples were swirled and inverted 2-3 times and settled for 2 minutes. The number of larvae floating was counted. 1 mL 20% sucrose was added to the conical, and samples were swirled, inverted, and settled. The number of floating larvae was recorded after each 1 mL addition. Additional sucrose was added, and floating larvae were counted until all larvae floated in each genotype. This experiment was repeated with 5 additional cohorts. The average sucrose concentration for all larvae to float in each genotype was compared using a paired t-test in Prism.

### Nile Red Staining

Late third instar larval brains from *adp*^*1*^ and *white berlin* were dissected in PBS and then fixed with 4% paraformaldehyde in PBS for 30 minutes^32^. The brains were washed with PBST three times for 20 minutes each before incubating overnight at 4°C in 1 ug/mL Nile Red in methanol. Finally, brains were washed twice with PBST for 30 minutes each before being mounted for imaging.

### Confocal Microscopy

All imaging was performed on a Zeiss LSM 710 confocal microscope with the 40X water immersion lens. A single brain lobe was centered in frame^20,21^. Zoom was set to 0.7, and scanning was done at speed 9. Z-stacks were set to encompass the entire z-range of the lobe, and the stack size was set at 2 μm.

#### Nile Red

The Rhodamine Red-X channel was used for Nile Red imaging. The frame size was 1024×1024 with a line averaging of 2.

#### Volume

Z stacks were set using Deadpan signal (647 channel) for volumetric analysis. The frame size was 512×512.

### Volume Analysis

Analysis was performed using the IMARIS software with the surface function. To compute volume, we drew surfaces around every 5th z-stack, including the two farthest ends of the brain. The stacks were then compiled, and volume was generated automatically. The average volume was compared using a t-test in Prism.

### Nile Red Analysis

Analysis was performed using the IMARIS software with the surface and spots functions. First, all images were set to the same brightness and contrast settings (Minimum of 35.46, maximum of 255, and gamma or 2.12). Volume was computed under the surface tab as described above, then a mask of the surface was generated to be used as a region of interest in spots. Under the spots tab, we used the automatic spots counter with an estimated XY diameter of 1.96523 units, and the quality filter was set to 20%, allowing for the most lipid droplets to be counted without generating false positives. For each brain, the number of lipid droplets was divided by the volume to compute the number of droplets/ μm^3^. The average number of lipid droplets/ μm^3^ was then compared between groups with a t-test in Prism.

### Generation of Humanized Flies

Wildtype *WDTC1* cDNA was obtained from the University of Utah Human ORF Core and is part of the Human ORF collection from the Mammalian Genome Collection^40^. The cDNA was cloned using Gateway cloning into pGW-HA.attB^41^ for expression under UASt regulatory sequences. The vector was injected into VK33 flies that contain an attP docking site at 65B2 on 3L, and transgenics were generated using phiC31 integrase-mediated recombination into the third chromosome.

### Statistical Analysis

All statistical analyses were performed using GraphPad Prism software. Independent t-tests for brain volume of neural stem cell knockdown, post-mitotic neuronal knockdown, Oregon-R vs adp60, lipid droplet analysis, and white berlin vs adp1 were done by selecting “t-tests” under column analyses and “Unpaired” under Experimental Design and assuming Gaussian distribution and equal SD. The paired t-test for WDTC1 rescue of buoyancy was performed by selecting “t-tests” under column analysis and “Paired” under Experimental Design and assuming Gaussian distribution and equal SD. Finally the repeated measures one-way ANOVA for the buoyancy in Figure 2C was performed by selecting “One-Way ANOVA” under Column Analyses, “each row represents matched, or repeated measures, data” under Experimental Design, assume Gaussian distribution, and not assuming sphericity therefore using Geisser-Greenhouse correction. Under the “Multiple Comparisons” tab, “Compare the mean of each column to the mean of every other column” to allow for identification of rescue phenotypes. The p-values reported in figure legends were the P value under the Repeated Measures ANOVA Summary and the adjusted P values from the multiple comparisons.

